# Mapping Social Relevance in Affective Scenes: A Large-Scale Multidimensional Rating Study

**DOI:** 10.1101/2025.08.14.670229

**Authors:** Vanessa Mitschke, Francesco Grassi, Alperen Doganer, Anne Schacht

**Affiliations:** Department for Cognition, Emotion and Behavior, Institute of Psychology, University of Goettingen, Goettingen, Germany; CRC 1528 Cognition of Interaction, Goettingen, Germany

**Author notes:** Corresponding Author: (AS).

**Keywords:** social relevance, affective scenes, stimulus norming, dimensional ratings, scene perception, social cognition

## Abstract

Understanding how individuals evaluate social content in affective scenes is crucial for research in emotion, social cognition, and decision-making. However, standardized image databases often lack fine-grained ratings of social dimensions beyond basic emotional content. We present a normative dataset of 296 affective scenes rated by 424 adult participants across six social-affective dimensions: social relevance, emotion sharing, action sharing, interpersonal equality, scene pleasantness, and participatory arousal. Each image was independently coded for objective features such as number of people, face visibility, and gaze direction.

All rating dimensions showed excellent interrater reliability. Pleasantness ratings demonstrated strong convergent validity with established valence norms from the International Affective Picture System (IAPS), while arousal ratings differentiated between experienced and participatory arousal, suggesting that the latter captures a distinct affective component. Correlation analyses revealed that social relevance was strongly predicted by scene pleasantness and a composite “engagement” factor combining emotion sharing, action sharing, and equality. Notably, the association between pleasantness and social relevance was especially pronounced in scenes depicting single individuals. Linear mixed models further indicated that extraversion modestly amplified the relationship between engagement and social relevance, while other personality traits and empathy did not significantly affect ratings.

This open-access dataset provides a reliable, multidimensional tool for selecting affective scenes based on both emotional and social-interpersonal features. It supports improved stimulus control and design in a wide range of behavioral and neurocognitive research. By systematically mapping social meaning in static affective scenes, our study advances existing methodological resources and offers empirical insight into the structure of social-affective appraisals. All image codes, mean ratings, and analysis scripts are publicly available via the Open Science Framework.

## Introduction

Detecting and interpreting social information from visual cues is crucial for navigating complex environments. Humans are adept at extracting affective and social meaning from visual scenes, often within milliseconds of exposure. Experiments involving the perception and evaluation of social information have been integral to our understanding of complex interactions between emotional, motivational, and cognitive processes. While research has extensively characterized emotional responses to visual scenes using databases such as the International Affective Picture System (IAPS; [1]), far less is known about how people perceive and evaluate the social qualities of these complex scenes, particularly when they depict interactions among multiple individuals.

Despite growing interest, there is no consensus on how to define or operationalize social information in complex scenes. Some studies rely on binary categories (e.g., presence of humans; [2]), while others focus on interpreting social cues [3] or assessing levels of social engagement [4]. In this study, we define social relevance as the perceived importance of a stimulus for understanding, predicting, or responding to others’ behavior. This includes cues related to interpersonal dynamics, intentions, and interactions [5,6]. Like emotional valence, social relevance must be inferred from contextual and relational information [7], making it a psychologically meaningful but methodologically challenging construct to quantify.

Research on social perception has largely concentrated on either isolated features such as faces, their emotional expressions, or gaze-driven attentional cues, often neglecting the complexity of real-world social interactions (for a detailed discussion; see). Such interactions are difficult to capture with isolated face stimuli; instead, they require scenes that depict full-body postures, multiple agents, and contextual (non)social cues. Recent studies suggest that social interaction influences attentional capture [9], and perceived pleasantness [10], beyond the simple presence of one or two humans.

This omission of interactional complexity is nontrivial. Social cues are not only a key driver of attention [6] but also modulate physiological and behavioral responses in affective tasks [11]. A growing body of research highlights the automatic attentional prioritization of social features such as faces, gaze direction, or interaction cues—even when emotional content is held constant [5,12–14]. These findings suggest that social relevance constitutes a distinct and powerful dimension of stimulus processing – interacting with, but not reducible to, emotional valence.

This perspective is increasingly supported by recent empirical findings from behavioral and neuroscientific paradigms. For example Kosonogov et al. [15] demonstrated that increasing the number of people in affective scenes systematically altered valence and arousal ratings, highlighting the influence of social content on affective evaluations. However, their approach treated social presence as a categorical feature and did not capture the nuances of interaction which are conveyed in social scenes. Our study extends this line of work by employing fine-grained, continuous ratings of key social-affective dimensions — including perceived social relevance, action and emotion sharing, and interpersonal equality — to better capture how social meaning is evaluated.

Additional evidence underscores the robustness and early impact of social relevance in complex scene processing. Social information can shift emotional priorities, alter visual salience, and guide attention—even under cognitive load [16]. For instance, emotional salience can override low-level visual features [17], and social relevance shapes both attentional dynamics and memory representations [3,18]. Neuroimaging findings further indicate that high-level social-affective features—rather than purely perceptual characteristics—primarily drive the neural encoding of observed actions [19]. Together, these findings reinforce the need for normative databases that include structured social-affective dimensions to better reflect real-world scene interpretation.

Structured neural representations further support the idea that social relevance is cognitively organized along interpretable dimensions. For example, Zhang et al. [20] showed that social decisions are encoded in a two-dimensional space of affiliation and power within hippocampal and parietal regions. Similarly, [21] found that the dorsomedial prefrontal cortex abstracts social interactions into basis functions that generalize across multi-agent contexts. These findings demonstrate that social meaning is systematically encoded and can be captured using standardized rating protocols.

A recent data-driven taxonomy by Santavirta et al. [22] provides further support. Using dynamic film clips, they identified both categorical and continuous dimensions of perceived social meaning, such as emotion sharing, power dynamics, and action coordination. Their results show that ecologically valid social features are consistently appraised across observers. This aligns closely with our approach, which aims to empirically map and validate key dimensions of social relevance in static visual scenes. Our study diverges from theirs by using affective static images— common in visual neuroscience—and by linking social dimensions to valence and arousal ratings. We further conduct exploratory factor analyses to uncover the latent structure of social-affective appraisals, offering a validated, open-access dataset and visualization tool for broader use.

### The present Study

Whereas previous research has often relied on binary classifications of social content or focused narrowly on facial features, our study offers a comprehensive, dimensional approach to mapping social meaning in affective scenes. By combining continuous ratings of key social-affective dimensions with objective image features and established valence/arousal norms, we provide a structured resource for selecting and analyzing social stimuli in experimental research. This multidimensional dataset enables researchers to move beyond coarse contrasts—such as social versus non-social images—and instead target specific interpersonal properties, such as emotion sharing, action coordination, and equality. In doing so, our work supports more precise stimulus control and contributes to a richer understanding of how individuals appraise social relevance in complex visual environments.

We conducted a large-scale rating study of 296 affective images, focusing on evaluations of social and interpersonal content. In addition to standard ratings of scene pleasantness and participatory arousal, participants provided judgments of emotion sharing, action sharing, and interpersonal equality. Objective features such as number of visible individuals, face visibility, and gaze direction were coded independently.

We adopted an exploratory approach to investigate social-affective processing in complex visual scenes. Rather than testing specific hypotheses, we pursued three primary objectives. First, we aimed to establish the reliability and construct validity of the rating dimensions, and to examine the relationship between perceived social features and valence. We assessed interrater agreement and tested convergent validity by comparing our pleasantness and participation ratings with IAPS norms [1]. Second, we explored whether individual differences in personality traits and empathy influenced social relevance ratings. Third, we examined how the number of individuals present in a scene affected ratings across dimensions—allowing us to differentiate between scenes with single agents and those depicting multi-agent social interactions.

## Methods

### Participants

Participants were recruited online using a convenience sample provided by MakeOpinion (https://www.makeopinion.com/) from 12/12/2023 to 11/01/2024. Of the 497 recruited participants, eight did not give their consent, and four did not complete the study. Additionally, 47 participants were excluded because they failed the attention checks, and 14 where further excluded because their responses indicated lack of attention to the questions (see the paragraph on data processing).

All analysis were run on the final sample of 424 participants, between 18 and 64 years of age (*M* = 41.47, *SD* = 12.42). 37.03% of the participants identified as female, 62.50% as male, 0.24% as transgender male, and 0.24% preferred not to declare. **Table 1** shows further details of the demographics of the final sample.

**Table 1.**
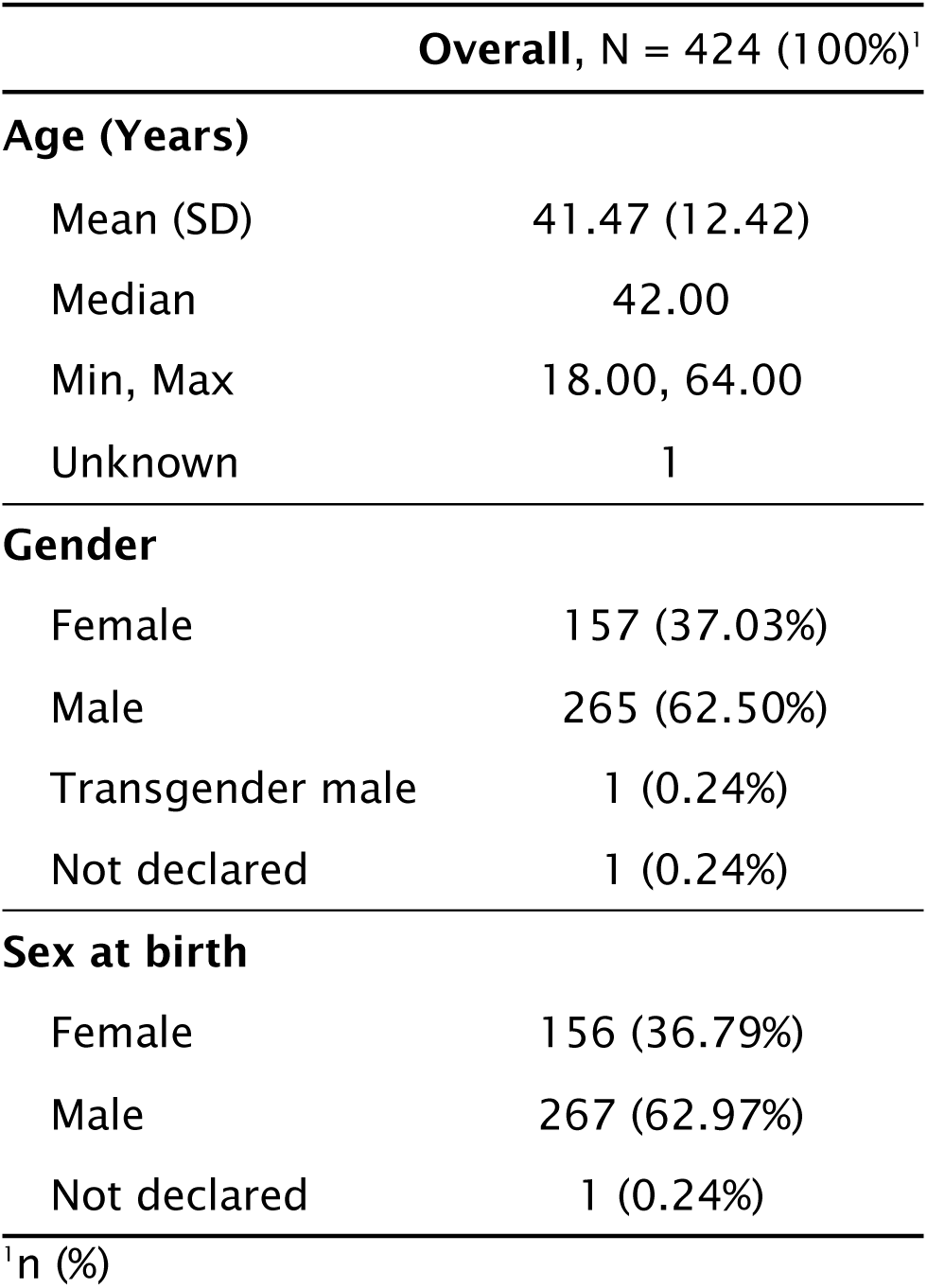
Demographics table of the final sample of participants.

### Ethics

This study was approved by the Ethics Committee of the Georg-Elias-Müller Institute for Psychology at the University of Göttingen (Approval No. 293, follow-up to Approval No. 199). All participants provided written informed consent electronically before participation. Consent was obtained via an online form integrated with the Prolific platform, where participants actively clicked to confirm consent after reading the information sheet and GDPR/data protection statement. Only participants aged 18 years or older were eligible. No minors were included in the study.

### Image Selection

For the purpose of this study, we selected images from the IAPS database [1], focusing on scenes that depict individuals either alone, in dyadic interactions, or as part of a group. The selected images primarily portray everyday scenarios such as shopping, walking down the street, waiting at a bus stop, or engaging in casual conversation. Additionally, the set includes some images representing more intense but commonly encountered situations as portrayed in media, such as scenes of war, plane crashes, traffic accidents, and interpersonal conflict. To ensure ecological validity and ethical applicability in future research, we excluded images depicting extreme positive (e.g., erotic content) and extreme negative (e.g., body mutilation) affective content. This approach was intended to maintain a focus on naturalistic social contexts. Based on these criteria, an initial pool of 424 images was compiled. Duplicates and overly repetitive themes (e.g., multiple images of dental visits or children) were removed to improve the distinctiveness of the stimuli. The final image set was refined to include a balanced sample of 100 images per interaction category (single individuals, dyads, and groups), distributed as equally as possible across negative, neutral, and positive affective contexts as defined by the normed database ratings of valence [1]. A complete list of IAPS codes and associated data for the final stimulus set is available on the Open Science Framework (https://osf.io/9m3ef/?view_only=082eceb798aa4cf88792657ca1f03cb1).

### Procedure

After providing consent, giving image examples (exemplary images were not part of the rating), participants viewed 50 images randomly selected from one of three possible subsets of 100 images (order randomized across participants) and rated each one on perceptual and affective dimensions using 9-point Likert scales (for an overview of the rating procedure see **Figure 1**). First, participants were asked to assess the overall social relevance of the scene by responding to the question: “*Please look at the scene as a whole. How social is this scene?*” (1 = "not at all social", 5 = "neither", 9 = "very social"). Participants were then asked to imagine themselves as part of the depicted scene and evaluate its affective qualities. They rated scene pleasantness (“*How pleasant do you think it is to be in the situation depicted*?”; 1 = "very pleasant", 5 = "neither", 9 = "very unpleasant") and scene participation (“*Does the scene involve a lot of movement and active participation, or a slower and calmer participation?*”; 1 = "very active", 5 = "neither", 9 = "very calm"). Next, they indicated the number of individuals in the image by selecting whether they saw one person or more than one (1 = "one", 2 = "more than one"). For scenes containing more than one individual, participants completed additional ratings on interaction-based dimensions. These assessed emotional sharing (“*Are the individuals in the scene sharing the same emotional experience?*”; 1 = "completely shared", 5 = "neither", 9 = "completely separate”), with instructions clarifying that, for example, two people laughing would indicate shared emotion, while one laughing and one crying would indicate separate emotional experiences. They also rated action sharing (“*Are the actions depicted in the scene performed collectively, or is each individual doing something on their own?*”; 1 = "joined action", 5 = "neither", 9 = "separate actions”), with examples such as two people carrying a sofa together (joint action) or performing unrelated tasks like reading and fishing (separate actions). Lastly, participants rated equality in the scene (“*Are the individuals in the scene equals, or is one person more in control than the other?*”; 1 = "fully equal", 5 = "neither", 9 = "one person in control”), with examples including differences due to social roles or context, such as a police officer making an arrest or a teacher instructing a student. All ratings were obtained without time constrictions. All items were originally given in German (see *Item List German* in the online repository materials for the original wording).

**Figure 1.**
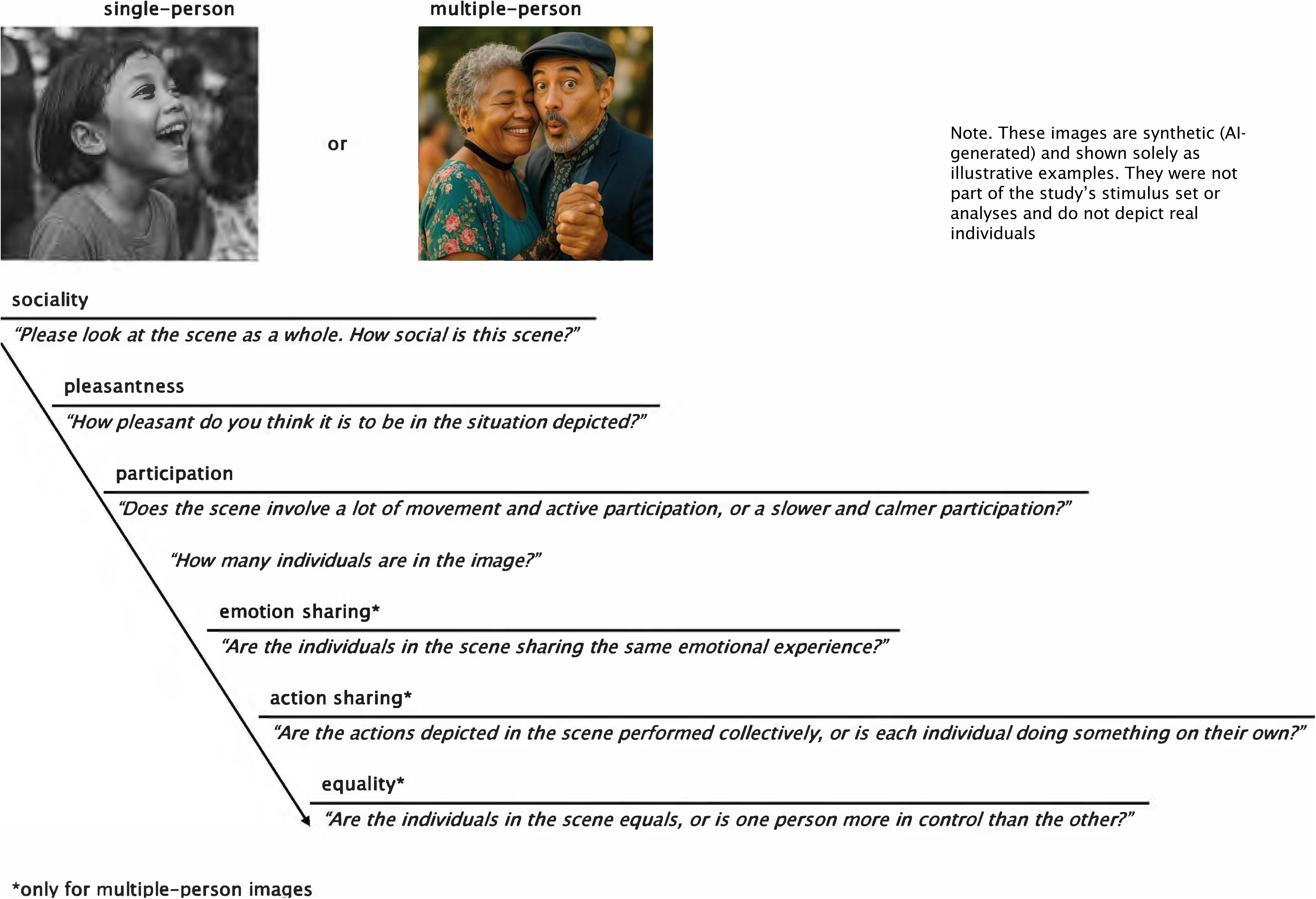
Trial scheme with questions and names of the respective rating variables used in the analysis. The images shown are illustrative examples from the OASIS database [25] and were not included since pictures used in the present study are copyrighted.

After the image ratings, all participants answered two questionnaires assessing personality and empathy traits. Personality traits were assessed using the Big Five Inventory (BFI-K; [23]) and empathy was assessed using the German version of the Interpersonal Reactivity Index (SPF; [24])

The objective image features were assessed by three independent coders and included the number of visible people and faces, grouping of people, visibility of faces, gaze sharing between people in the image, and eye contact with the viewer. The responses from the three coders were compared, and discrepancies were resolved by taking the majority response for each image and feature. Objective image features were then combined with participant responses.

### Data Processing

Processing of the data was conducted in R (version 4.5.0; [26]), using *tidyverse* (v2.0.0; [27]) and *readxl* (v1.4.3; [28]) packages. The first step of data processing involved excluding non-consenting participants, those who did not complete the study, and participants who failed to correctly respond to both attention checks. Participant responses regarding the social relevance of the image, scene pleasantness, scene participation, presence of single or multiple persons in the image, as well as emotion sharing, action sharing, and equality within the image were then extracted for further processing.

Since the study included additional questions when participants indicated that multiple people were depicted in an image, it was possible that some participants might have intentionally indicated that multiple-person images contained a single person to complete the study faster. To address this, participants who disagreed with the independent coders regarding the number of people in the image in more than 20% of the trials were excluded (*N* = 14). Moreover, as the number of people in the image was treated as an objective feature based on the rating of the independent coders, all trials where participants disagreed with the coders’ assessments were removed. This resulted in a loss of 4.41% of the trials.

The ratings for social relevance, scene pleasantness, scene participation, emotion sharing, action sharing, and equality were adjusted so that a value of 1 represents the lowest rating and a value of 9 represents the highest rating for each variable.

Participant scores on the SPF (empathy) questionnaire were calculated by first reversing the scoring on the negatively worded items and then averaging the scores of the items belonging to each of the questionnaire sub-scales (“-“ indicate reverse-scored items): *perspective taking* (items: 3-, 8, 11, 15-), *fantasy* (1, 5, 7-, 12-, 16), *empathic concern* (2, 4-, 9, 14-) and *personal distress* (6, 10, 13-). The same approach was used for the participant scoring on the BFI-K questionnaire, aggregating the scoring belonging to the sub-scales: *extraversion* (1-, 6, 11-, 16), *agreeableness* (2-, 7, 12-, 17-), *conscientiousness* (3, 8-, 13, 18-), *neuroticism* (4, 9-, 14, 19) and *openness* (5, 10, 15, 20, 21-).

### Interrater Reliability of Image Ratings

Interrater reliability of the subjective ratings was assessed using intraclass correlation coefficients (ICCs). Given that our design was partially crossed—each image was rated by a subset of participants (with participants rating only 50 of 100 images within one of three distinct image sets)—the appropriate approach was to calculate ICC(1, k), i.e., based on a mean-rating, absolute-agreement, one-way random effects model [29]:

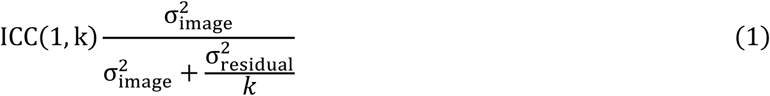

Where 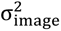 is the variance attributed to differences between images, 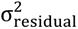 is the residual variance, and 𝑘 is the number of raters per image. This was done via the function *lmer* of the R package *lme4* (v1.1-37, [30]). Confidence intervals for the ICC estimates were obtained through parametric bootstrapping using the *bootMer* function (N = 10000 bootstraps).

### Convergent Validity with IAPS Scales

To assess potential convergent validity between our rating scales and the norms of the IAPS database, Spearman’s correlation coefficients were calculated for each rating variable (social relevance, scene pleasantness, scene participation, emotion sharing, action sharing, equality) with both the valence and arousal norms of the IAPS database.

### Exploratory Correlation Analysis

Correlation analysis was conducted to explore relationships between image ratings and objective image features. Since these variables are of different scales (i.e., interval, ordinal, nominal), we used an approach based on so-called *difference semimatrices* proposed by Podani et al. (2023). Briefly, given a dataset with 𝑛 variables (rows) and 𝑚 observations (columns), for each variable ℎ a 𝑚 × 𝑚 matrix 𝐷^(h)^ is calculated where each (𝑗, 𝑘) entry 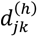 quantifies the difference between observations 𝑗 and 𝑘 on variable ℎ. The way these differences are calculated depends on the scale of the variable ℎ (see Podani et al. [31] for details). In a second step, the correlation between two variables ℎ and 𝑖 (referred to as *d-correlation* to distinguish it from the regular Pearson correlation) is calculated from the two semimatrices 𝐷^(h)^ and 𝐷^(i)^ as:

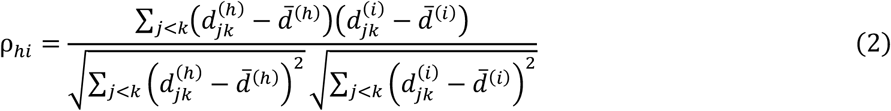

The analysis was conducted on the mean image ratings (averaged across participants) separately for single- and multiple-person images, since ratings of emotion sharing, action sharing and equality were available only for the latter.

We further visualized the average image ratings of scene pleasantness against scene participation to explore whether the two variables showed the typical “U-shaped” relationship of the IAPS norms of valence and arousal (**[Figure 3 about here]** Figure 3).

### Calculation of Composite *Engagement* Rating

The correlation analysis showed an almost perfect association between ratings of emotion sharing, action sharing and equality (see **Results** section), and an additional factor analysis confirmed very high loadings of the three variables on one latent factor (Supplementary Material **S1**). To mitigate multicollinearity concerns when incorporating these variables into subsequent analyses, we computed a composite measure. This was achieved by averaging each participant’s ratings of emotion sharing, action sharing, and equality for each image. The resulting composite measure, referred to here as *engagement* rating, was utilized in the following analysis steps.

### Analysis of the Effect of Scene Pleasantness on Social Relevance

A Linear Mixed Model (LMM;) was used to investigate the effect of scene pleasantness on ratings of social relevance. The model also controlled for potential effects of scene participation and gaze direction in respect to the viewer (i.e., direct or averted). These two terms were included as main fixed effects, as the exploratory analysis did not reveal any particular linear or non-linear relationship between these variables and scene pleasantness. Moreover, since the ratings of social relevance, scene pleasantness and scene participation showed different distributions for single- and multiple-person images (**Figure 2A-C**), an additional factor *person number* (with levels: single, multiple) was added in interaction with both scene pleasantness and scene participation.

**Figure 2.**
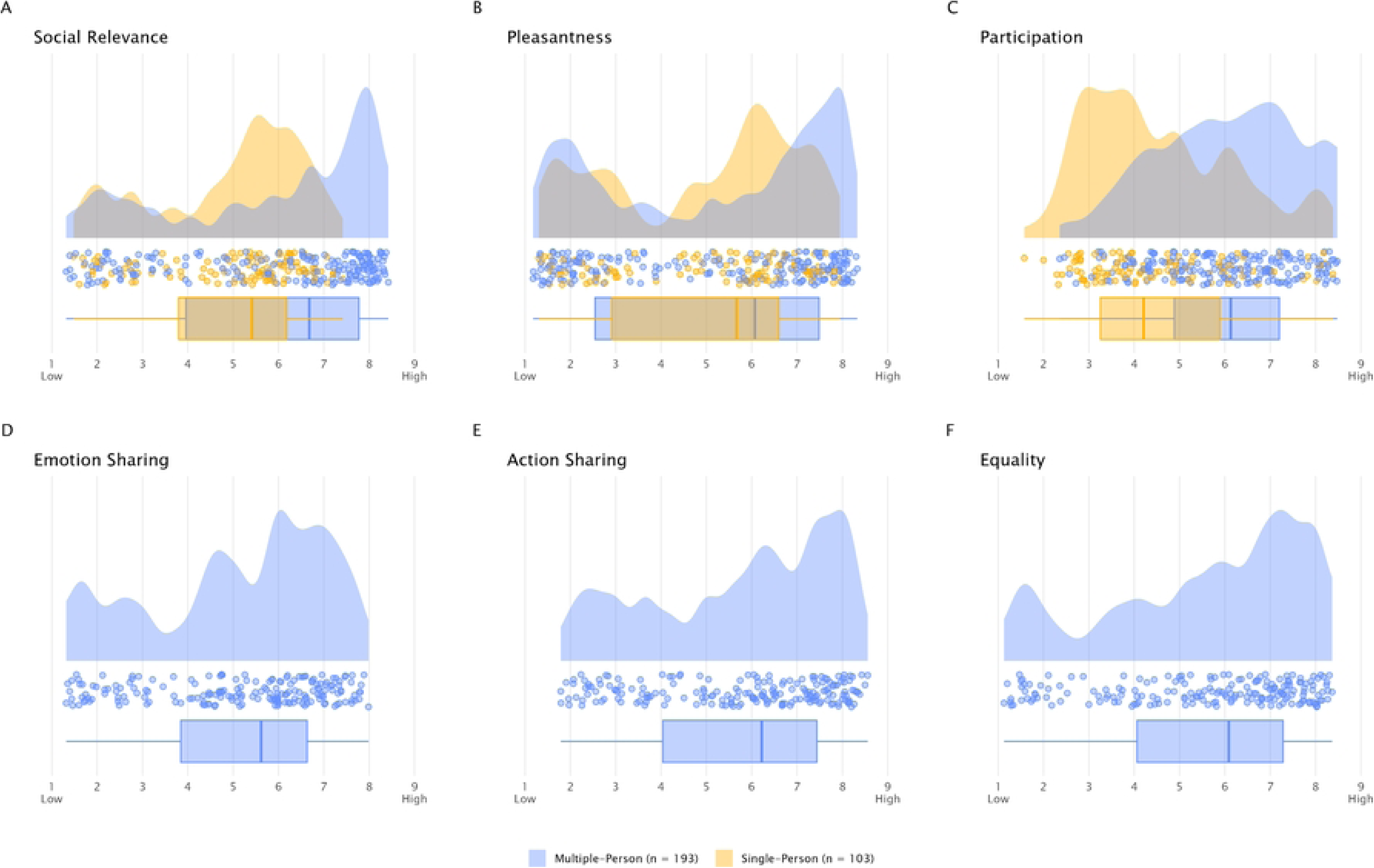
Raincloud plots showing the distribution of mean ratings across the images included in the study. Each dot represents an image’s mean rating (averaged across participants), the central box indicates the interquartile range with the median bar, and the density portion illustrates the overall shape of the distribution

Participant ID and image ID were included as random intercepts. To maintain a 5% Type I error rate, all identifiable random slopes and their correlation with random intercepts were included [33,34], specifically for scene pleasantness and scene participation within participant ID. The full model therefore was defined as:

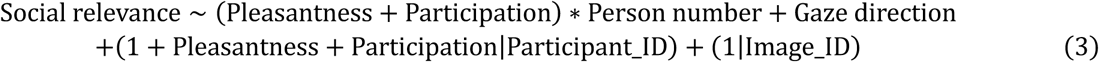

The overall effect of scene pleasantness was tested using a full-null model comparison [35], where the null model lacked the fixed effect of scene pleasantness but was otherwise identical to the full model.

### Analysis of the Effect of Engagement on Social Relevance

Another LMM was used to investigate the effect of the composite rating of engagement on the social relevance of the images. Even in this case the model controlled for the additional effects of scene participation and gaze direction. However, since the rating of engagement was only available for multiple-person images, no further effect of person number was investigated. Participant ID and image ID were included as a random intercept, and engagement and scene participation within participant ID as random slopes. The full model was therefore defined as:

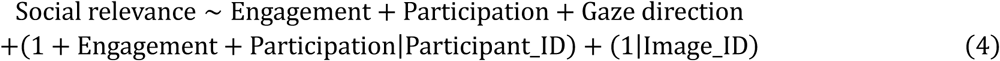

The corresponding null model lacked the fixed effect of engagement but was otherwise identical to the full model.

### Analysis of the Moderation Effects of Personality Traits

We further investigated whether participants personality traits could moderate the effects of scene pleasantness and engagement on social relevance. To this aim, two additional LMMs were defined, including the interaction effects of each sub-scale of the SPF and BFI-K with scene pleasantness or engagement, respectively. The moderation effect of personality traits was tested by comparing each model with a reduced one lacking the interaction term while being otherwise identical to the respective full model.

Even in this case, main effects of scene participation and gaze direction were included as control predictors, participant ID and image ID as random intercept, and scene pleasantness/engagement and scene participation within participant ID as random slopes.

All LMMs described above were fitted in R with the function *lmer* from the *lme4* package. Before fitting the models, all numerical predictors were z-transformed (mean-centered and scaled) to provide a valid reference level. The significance of the fixed effects was tested using the Satterthwaite approximation [36] via the function *lmer* from the *lmerTest* package (version 3.1- 3; [37]). The assumptions of normally distributed and homogeneous residuals were checked by visually inspecting QQ-plots of the residuals [38] and by plotting residuals against fitted values [39]. Model stability at the level of estimated coefficients and standard deviation was assessed by excluding one level of the random effect (i.e., participant ID and image ID) at a time and refitting the model [40], using a function kindly provided by Roger Mundry (University of Göttingen). Confidence intervals for the model estimates were obtained by parametric bootstrapping using the *bootMer* function from the *lme4* package, with 1000 bootstraps. Due to the different computation methods, the interpretation of Satterthwaite approximated p-values and bootstrapped CIs might differ. The absence of collinearity issues was verified by calculating the Variance Inflation Factors (VIF) for a model without interaction terms, using the *vif* function from the *car* package (version 3.1-3; [41]).

## Results

Descriptive statistics of all the image rating variables and image objective features is presented in **Table 2**, while **Figure 2** shows the distribution of the image rating variables.

**Table 2.** Descriptive statistics of the image rating variables and objective features.

| Overall, N = 296 (100%)<sup>1</sup> |  |
| --- | --- |
| **Social Relevance** |  |
| Mean (SD) | 5.54 (2.09) |
| Median | 5.94 |
| Min, Max | 1.32, 8.42 |
| **Pleasantness** |  |
| Mean (SD) | 5.13 (2.33) |
| Median | 5.86 |
| Min, Max | 1.18, 8.33 |
| **Participation** |  |
| Mean (SD) | 5.50 (1.70) |
| Median | 5.60 |
| Min, Max | 1.58, 8.48 |
| **Emotion Sharing** |  |
| Mean (SD) | 5.13 (1.90) |
| Median | 5.62 |
| Min, Max | 1.32, 7.99 |
| **Action Sharing<sup>2</sup>** |  |
| Mean (SD) | 5.75 (1.99) |
| Median | 6.22 |
| Min, Max | 1.79, 8.56 |
| **Equality<sup>2</sup>** |  |
| Mean (SD) | 5.55 (2.11) |
| Median | 6.08 |
| Min, Max | 1.13, 8.37 |
| **Face Visibility** |  |
| All | 157 (53.04%) |
| Partial | 102 (34.46%) |

| **Overall, N = 296 (100%)<sup>1</sup>** |  |
| --- | --- |
| None | 37 (12.50%) |
| **Gaze Direction Viewer** |  |
| Direct | 80 (27.03%) |
| Averted | 216 (72.97%) |
| **Gaze Direction Within** |  |
| Yes | 35 (11.82%) |
| No | 229 (77.36%) |
| Unclear | 32 (10.81%) |
| **N Face** |  |
| 0 | 37 (12.50%) |
| 1 | 161 (54.39%) |
| 2 | 55 (18.58%) |
| 3 | 20 (6.76%) |
| 4 | 11 (3.72%) |
| 5 | 4 (1.35%) |
| 6<sup>3</sup> | 8 (2.70%) |
| **N People** |  |
| 1 | 122 (41.22%) |
| 2 | 100 (33.78%) |
| 3 | 24 (8.11%) |
| 4 | 15 (5.07%) |
| 5 | 6 (2.03%) |
| 6<sup>3</sup> | 29 (9.80%) |
| **People Group** |  |
| Single | 103 (34.80%) |
| Dyad | 94 (31.76%) |
| Group | 99 (33.45%) |

<sup>1</sup>n (%)
<sup>2</sup>Only multiple-person images (N = 193)
<sup>3</sup>6 or more

### Interrater Reliability of Image Ratings

ICC estimates and their 95% confidence intervals revealed excellent [29] interrater reliability for all image rating variables (**Table 3**.).

**Table 3.** ICC estimates and 95% confidence intervals of the image rating variables.

| Rating Variable | ICC | 95% CI (Lower - Upper) | N (images) | Avg. Raters per Image |
| --- | --- | --- | --- | --- |
| Social Relevance | 0.99 | 0.99 - 0.99 | 296 | 68.46 |
| Pleasantness | 0.99 | 0.99 - 0.99 | 296 | 68.46 |
| Participation | 0.98 | 0.97 - 0.98 | 296 | 68.46 |
| Emotion Sharing | 0.98 | 0.98 - 0.99 | 193 | 67.28 |
| Action Sharing | 0.98 | 0.98 - 0.99 | 193 | 67.28 |
| Equality | 0.99 | 0.98 - 0.99 | 193 | 67.28 |

### Convergent Validity with IAPS normative data (valence and arousal)

**Table *4*** shows the results of the Spearman’s correlation of each of the rating variables and the IAPS norms of valence and arousal. All rating variables, except scene participation in multiple-person images, presented a statistically significant correlation with valence norms, with a very strong positive association for social relevance and scene pleasantness, a strong positive association for emotion sharing, action sharing and equality and a weak association for scene participation in single-person images. Conversely, all rating variables showed a statistically significant correlation with arousal scores, moderately positive for scene participation, moderately negative for social relevance and pleasantness, and weakly negative otherwise.

**Table 4.** Spearman’s correlations of rating variables and IAPS norms of valence and arousal.

| Rating Variable | IAPS |  |  |  |  |  |
| --- | --- | --- | --- | --- | --- | --- |
|  | Valence |  |  | Arousal |  |  |
|  | rho | df | p | rho | df | p |
| **Single-Person images** |  |  |  |  |  |  |
| Social Relevance | 0.85 | 103 | < .001 | -0.36 | 103 | < .001 |
| Pleasantness | 0.90 | 103 | < .001 | -0.33 | 103 | .001 |
| Participation | 0.34 | 103 | < .001 | 0.41 | 103 | < .001 |
| **Multiple-Person images** |  |  |  |  |  |  |
| Social Relevance | 0.87 | 193 | < .001 | -0.49 | 193 | < .001 |
| Pleasantness | 0.91 | 193 | < .001 | -0.52 | 193 | < .001 |
| Participation | -0.06 | 193 | .415 | 0.58 | 193 | < .001 |
| Emotion Sharing | 0.73 | 193 | < .001 | -0.32 | 193 | < .001 |
| Action Sharing | 0.67 | 193 | < .001 | -0.27 | 193 | < .001 |
| Equality | 0.62 | 193 | < .001 | -0.38 | 193 | < .001 |

### Exploratory Correlation Analysis

The full results of the exploratory correlation analysis are presented in **Table S1** and **Table S2** in the Supplementary Material).

Three patterns in particular emerged from the analysis. First, social relevance showed a very strong positive correlation with scene pleasantness (single-person images: *d* = .95, multiple-person images: *d* = .92) and a strong association with emotion sharing, action sharing and equality (*d* = .78, .76, .70, respectively).

Second, pleasantness and participation showed only a week association for single-person images (*d* =.31) and essentially no association for multiple-person images (*d* = -.04). The absence of a linear association does not necessary exclude some other forms of quadratic or non-linear relationship between these two variables, as in the case of the typical “U-shape” of the IAPS valence vs arousal. However, as can be seen in **Figure 3A**, no clear association, linear or otherwise, can be observed between the two variables.

**Figure 3.**
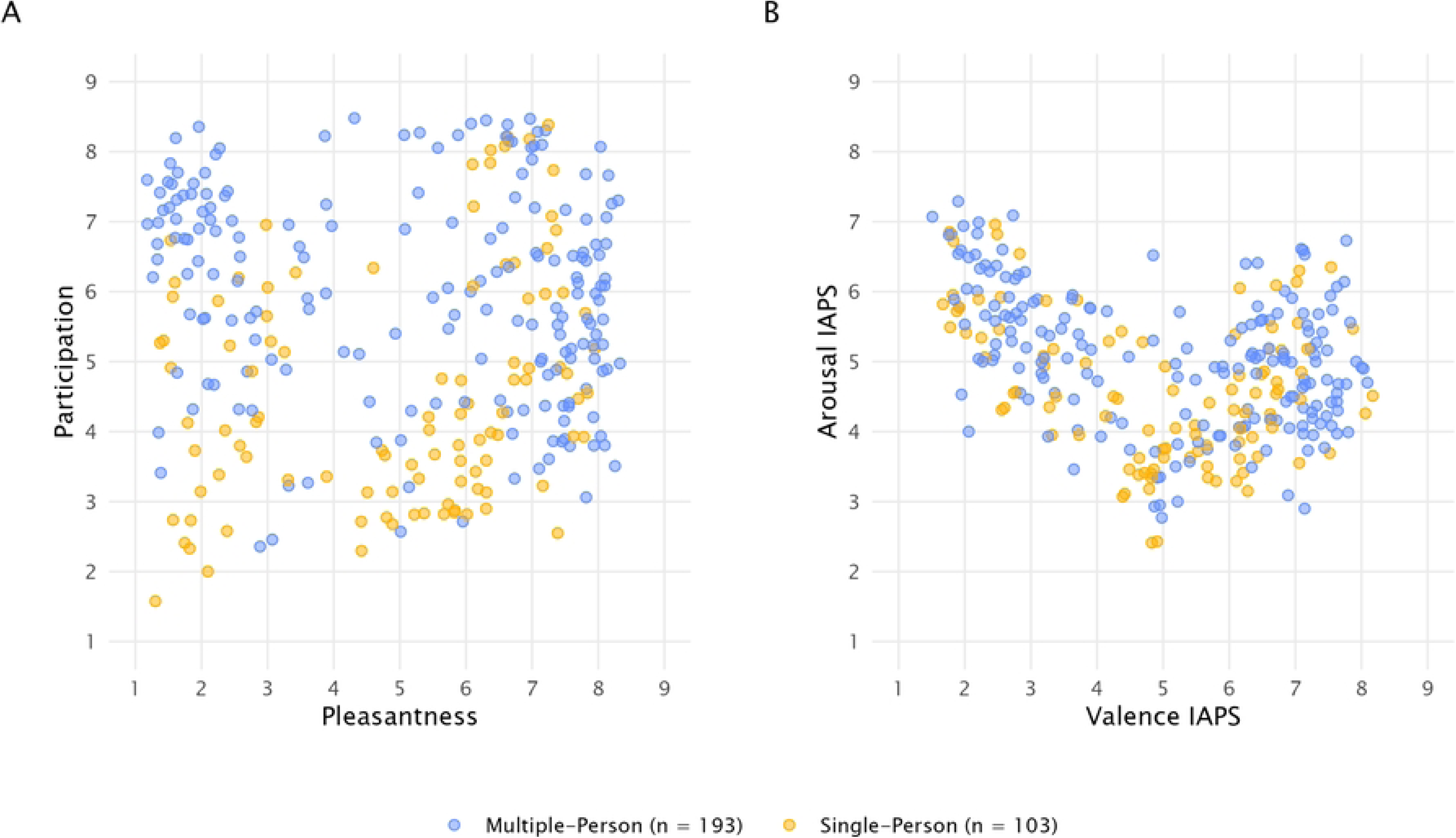
(A) Average image ratings of pleasantness vs participation. (B) IAPS norm of valence vs arousal for the same images used in the current study.

Third, emotion sharing, action sharing and equality showed strong or very strong positive associations among each other (emotion sharing vs action sharing: *d* = .95; emotion sharing vs equality: *d* = .90; action sharing vs equality: *d* = .81), suggesting that these three variables might tap one latent factor which can be interpreted as a measure of *engagement* in the image (see **Supplementary Material S2** for an additional exploratory factor analysis further supporting this interpretation).

All LMMs used in the analyses showed no multicollinearity issue and good stability. Visual inspection of the model diagnostics indicated that the assumptions of normality and homogeneity were reasonably met.

### Effect of Scene Pleasantness on Social Relevance

The effect of scene pleasantness on the ratings of social relevance was tested while controlling for scene participation, gaze direction and number of persons in the scene. The analysis showed that scene pleasantness significantly influences social relevance (full-null model comparison: *χ²* = 880.04, *df* = 15, *p* < .001). More specifically, the model showed a significant interaction of scene pleasantness and person number (*β* = -0.10, *SE* = 0.04, *p* = .006, 95% CI [- 0.16, -0.03], **Table 5**). Increased ratings of scene pleasantness resulted in increased social relevance, and this association was slightly stronger for single-person images compared to multiple-person ones (**Figure 4A**).

**Figure 4.**
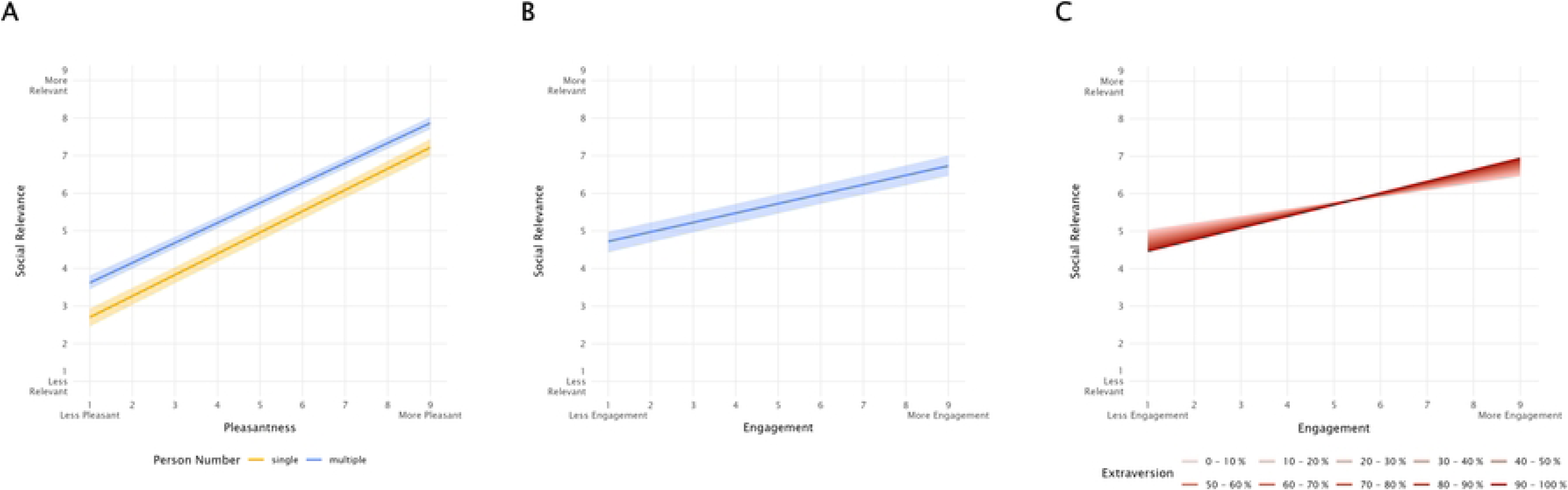
Social relevance as a function of (A) scene pleasantness, for single- and multiple-person images, (B) engagement, (C) engagement and extraversion. Shaded areas represent 95% confidence intervals.

**Table 5.** Summary of the LMM regarding the effect of scene pleasantness on social relevance.

| Predictors | R<sup>2</sup> | Estimate | std. Error | 95% CI | t value | p |
| --- | --- | --- | --- | --- | --- | --- |
|  | .43 |  |  |  |  |  |
| intercept |  | 5.23 | 0.12 | 4.99 5.47 |  | (1) |
| pleasantness<sup>(2)</sup> |  | 1.63 | 0.04 | 1.55 1.71 | 41.37 | < .001 |
| participation<sup>(2)</sup> |  | 0.21 | 0.03 | 0.16 0.26 | 7.80 | < .001 |
| person number (multiple)<sup>(3)</sup> |  | 0.78 | 0.12 | 0.55 1.00 | 6.49 | < .001 |
| gaze direction (averted)<sup>(4)</sup> |  | -0.25 | 0.13 | -0.50 0.01 | -1.96 | .052 |
| pleasantness<sup>(2)</sup> * person number (multiple)<sup>(3)</sup> |  | -0.10 | 0.04 | -0.16 -0.03 | -2.72 | .006 |
| participation<sup>(2)</sup> * person number (multiple)<sup>(3)</sup> |  | -0.16 | 0.03 | -0.22 -0.11 | -5.75 | < .001 |

(1) not shown because of being of very limited interpretability
(2) z-transformed to mean of zero and a standard deviation of one
(3) comparison with the reference level (single)
(4) comparison with the reference level (direct)

### Effect of Engagement on Social Relevance

Our second model tested the effect of engagement on social relevance, controlling for scene participation and gaze direction. The test revealed a significant influence of engagement on social relevance (full-null model comparison: *χ²* = 340.94, *df* = 12, *p* < .001). As can be seen in **Figure 4B**, increased engagement resulted in increased ratings of social relevance (*β* = 0.63, *SE* = 0.03, *p* < .001, 95% CI [0.58, 0.69], **Table 6**).

**Table 6.** Summary of the LMM regarding the effect of engagement on social relevance.

| Predictors | R2 | Estimate | std. Error | 95% CI | t value | p |
| --- | --- | --- | --- | --- | --- | --- |
|  | .08 |  |  |  |  |  |
| intercept |  | 6.55 | 0.31 | 5.90 7.19 |  | (1) |
| engagement<sup>(2)</sup> |  | 0.63 | 0.03 | 0.58 0.69 | 21.34 | < .001 |
| participation<sup>(2)</sup> |  | 0.10 | 0.03 | 0.05 0.15 | 3.92 | < .001 |
| gaze direction (averted)<sup>(3)</sup> |  | -0.84 | 0.35 | -1.54 -0.11 | -2.43 | .017 |

(1) not shown because of being of very limited interpretability
(2) z-transformed to mean of zero and a standard deviation of one
(3) comparison with the reference level (direct)

### Moderation of Scene Pleasantness Effects by Personality Traits

Participants’ personality traits were tested as potential moderators of the effect of scene pleasantness on social relevance. This was done by fitting a full model including the interaction effects of each SPF and BFI-K sub-scale and scene pleasantness, and comparing it to a reduced model lacking such interaction.

The comparison showed no significant difference between the full and the reduced model (*χ²* = 11.26, *df* = 9, *p* ≥ .05), suggesting that personality traits play no role in moderating the effect of scene pleasantness on ratings of social relevance.

The absence of a moderation effect does not imply that personality traits alone might not have a direct effect on social relevance. To test this, the reduced model was compared to a null model lacking the SPF and BFI-K sub-scales predictors entirely. This comparison was also not significant (*χ²* = 16.07, *df* = 9, *p* ≥ .05), indicating that, when accounting for the role of scene pleasantness (as well as controlling for scene participation and gaze direction) personality traits do not affect ratings of social relevance, nor directly, nor acting as moderators.

### Moderation of Engagement Effects by Personality Traits

Personality traits were also tested as potential moderators of the effects of engagement on social relevance. In this case, the full-reduced model comparison was significant (*χ²* = 20.95, *df* = 9, *p* = .013), indicating that, as a whole, personality traits moderated the relationship between engagement and social relevance. More precisely, the analysis of the individual fixed effects showed a significant interaction of engagement and extraversion (*β* = 0.06, *SE* = 0.02, *p* < .012, 95% CI [0.01, 0.1],

**Table *7***). Consistent with the results reported above, increased engagement resulted in increased ratings of social relevance, and this effect was stronger the higher the participant’s level of extraversion (**Figure 4C**).

**Table 7.** Summary of the LMM regarding the moderation by personality traits on the effects of engagement on social relevance.

| Predictors | R2 | Estimate | std. Error | 95% CI | t value | p |
| --- | --- | --- | --- | --- | --- | --- |
|  | .09 |  |  |  |  |  |
| intercept |  | 6.55 | 0.32 | 5.96 7.16 |  | (1) |
| engagement<sup>(2)</sup> |  | 0.63 | 0.03 | 0.57 0.68 | 21.58 | < .001 |
| empathic concern<sup>(2)</sup> |  | -0.00 | 0.03 | -0.07 0.06 | -0.08 | .940 |
| fantasy<sup>(2)</sup> |  | 0.03 | 0.04 | -0.05 0.10 | 0.69 | .493 |
| personal distress<sup>(2)</sup> |  | 0.01 | 0.04 | -0.06 0.08 | 0.26 | .791 |
| perspective taking<sup>(2)</sup> |  | -0.00 | 0.04 | -0.07 0.06 | -0.13 | .896 |
| agreeableness<sup>(2)</sup> |  | -0.00 | 0.04 | -0.08 0.08 | -0.05 | .955 |
| conscientiousness<sup>(2)</sup> |  | 0.01 | 0.04 | -0.07 0.07 | 0.16 | .875 |
| extraversion<sup>(2)</sup> |  | 0.00 | 0.04 | -0.08 0.08 | 0.12 | .906 |
| neuroticism<sup>(2)</sup> |  | -0.01 | 0.04 | -0.08 0.07 | -0.23 | .816 |
| openness<sup>(2)</sup> |  | 0.18 | 0.04 | 0.12 0.26 | 4.97 | < .001 |
| participation<sup>(2)</sup> |  | 0.10 | 0.03 | 0.05 0.15 | 3.92 | < .001 |
| gaze direction (averted)<sup>(3)</sup> |  | -0.84 | 0.35 | -1.49 -0.17 | -2.43 | .017 |
| engagement<sup>(2)</sup> * empathic concern<sup>(2)</sup> |  | -0.02 | 0.02 | -0.06 0.02 | -0.89 | .381 |
| engagement<sup>(2)</sup> * fantasy<sup>(2)</sup> |  | 0.03 | 0.02 | -0.02 0.07 | 1.11 | .274 |
| engagement<sup>(2)</sup> * personal distress<sup>(2)</sup> |  | 0.03 | 0.02 | -0.01 0.08 | 1.40 | .171 |
| engagement<sup>(2)</sup> * perspective taking<sup>(2)</sup> |  | -0.03 | 0.02 | -0.08 0.02 | -1.33 | .189 |
| engagement<sup>(2)</sup> * agreeableness<sup>(2)</sup> |  | 0.03 | 0.02 | -0.02 0.07 | 1.22 | .230 |
| engagement<sup>(2)</sup> * conscientiousness<sup>(2)</sup> |  | -0.01 | 0.02 | -0.05 0.03 | -0.44 | .664 |
| engagement<sup>(2)</sup> * extraversion<sup>(2)</sup> |  | 0.06 | 0.02 | 0.01 0.11 | 2.57 | .012 |
| engagement<sup>(2)</sup> * neuroticism<sup>(2)</sup> |  | 0.02 | 0.02 | -0.03 0.07 | 0.77 | .450 |
| engagement<sup>(2)</sup> * openness<sup>(2)</sup> |  | 0.02 | 0.02 | -0.03 0.06 | 0.69 | .492 |

(1) not shown because of being of very limited interpretability
(2) z-transformed to mean of zero and a standard deviation of one
(3) comparison with the reference level (direct)

## Discussion

This study set out to systematically map how individuals appraise social relevance in affective scenes by collecting continuous ratings along key social-affective dimensions and linking them to both objective image features and affective experience. We found that all rating dimensions showed excellent interrater reliability—indicating that participants applied the scales consistently and interpreted the constructs in a shared, reliable manner—and strong convergent validity with existing affective norms, supporting the robustness of the social-affective appraisal framework. Critically, social relevance was best predicted by scene pleasantness and a composite measure of interpersonal engagement—reflecting shared emotion, joint action, and perceived equality—highlighting that social meaning in complex scenes is both reliably perceived and structured along interpretable dimensions.

All rating dimensions showed excellent interrater reliability, confirming that participants applied the scales consistently and interpreted the constructs similarly. Pleasantness ratings showed strong convergent validity with the original IAPS valence norms, indicating a close correspondence between experienced valence during passive viewing and evaluations of pleasantness when imagining oneself within the scene. In contrast, participation ratings diverged from IAPS norms, likely reflecting the distinct nature of our participatory arousal measure, which captures imagined engagement (e.g., active vs. calm participation) rather than stimulus-driven physiological arousal. This distinction suggests that participatory arousal may tap into a different facet of affective processing—one that is more closely tied to imagined involvement and action readiness than to passive emotional intensity.

Social relevance was most strongly predicted by scene pleasantness and by the degree of interpersonal engagement in the scene, as reflected in shared emotions, actions, and perceived equality. This suggests that social scenes are often perceived as more meaningful when they are also emotionally positive and interactive—capturing reciprocal exchanges between individuals. Interestingly, the association between pleasantness and social relevance was strongest in scenes with limited social context (i.e., images depicting a single individual), indicating that even subtle interpersonal cues may activate social appraisal mechanisms when interpreted as affiliative or inviting. The relationship between social relevance and pleasantness was not significantly moderated by most personality traits or empathy, pointing to a partially universal structure of social-affective evaluation across perceivers. The only exception was extraversion, which amplified the link between engagement and social relevance. This may reflect a greater attunement among extraverted individuals to opportunities for social connection, consistent with trait-level differences in sociability and reward sensitivity in social contexts [42,43].

Our findings offer a methodologically grounded contribution to the emerging effort to formalize the construct of social relevance in visual scene perception. In this context, our study complements and extends the recent work of [22], who proposed a data-driven taxonomy of social features based on observer ratings of dynamic film clips. Their findings revealed that dimensions such as emotion sharing, power asymmetries, and action coordination are reliably perceived and cluster into meaningful patterns – underscoring the value of systematic feature annotation in understanding social perception. Building on this rationale, our study applies a similar dimensional framework to static affective images, which remain the standard format in many experimental paradigms across cognitive and affective neuroscience. This methodological focus not only enhances compatibility with existing databases like the IAPS but also supports wider adoption of structured social-affective ratings in stimulus selection and experimental design.

The strong correlation we observed between social relevance and pleasantness in single-person images suggests that even minimal interpersonal cues can influence evaluative appraisals. This pattern is consistent with appraisal theories, particularly the Component Process Model [44,45], which posit that early relevance checks—such as novelty, intrinsic pleasantness, and goal conduciveness – are sensitive to social contextual features. Our findings provide a concrete operationalization of these early appraisal components, showing how relational cues embedded in affective scenes systematically influence perceived social meaning. In this sense, our dimensional ratings capture not just descriptive scene features but also the psychological mechanisms through which individuals interpret and evaluate complex social contexts.

The strong association between perceived pleasantness and social relevance likely reflects the fundamental human need for belonging [46], which not only motivates social behavior but also shapes perceptual and evaluative processes. When individuals identify potential opportunities for interaction—particularly affiliative ones—these scenes are appraised as both socially meaningful and emotionally positive. This interpretation aligns with prior findings demonstrating that social cues can shift emotional processing priorities in complex scenes [3], and continue to guide attention even under cognitive load [18]. Such robustness is echoed in our own finding of high interrater agreement across social-affective dimensions, despite the complexity and variability of the scenes.

At the neural level, recent work has shown that social meaning is represented in structured, multidimensional formats. Zhang et al. [20] identified a network—including hippocampus and precuneus—that encodes individuals’ positions within a social space defined by power and affiliation. Similarly, Wittmann et al. [21] demonstrated that dorsomedial prefrontal cortex abstracts social interactions into compressed basis functions, enabling efficient representation of multi-agent contexts. Our study complements these findings by showing that static images—rather than interactive or dynamic tasks—can elicit reliably structured appraisals of social relevance. Moreover, we provide a validated tool to quantify these appraisals, facilitating systematic investigation of social relevance across diverse visual paradigms.

The fit between social relevance and affective pleasantness was not meaningfully influenced by individual differences in personality traits or empathy. Our results suggest that, as long as affective properties such as pleasantness and participation are accounted for, dispositional factors play a limited role in shaping social appraisals of visual scenes. This finding points toward a relatively universal and robust evaluation process, wherein observers consistently detect and respond to social cues—such as shared emotion or interpersonal equality—regardless of their baseline level of sociability, emotional sensitivity, or perspective-taking. Such consistency supports the view that certain social-affective appraisals may operate as early, automatic processes, sensitive to contextual features of the scene rather than stable internal traits. From a theoretical standpoint, this aligns with appraisal models proposing that relevance detection (e.g., goal conduciveness or intrinsic pleasantness) constitutes a fast, low-threshold mechanism that guides attention and meaning-making across individuals [44,45]. While more nuanced individual differences may still influence downstream emotional responses or decision-making, our findings suggest that the initial appraisal of social relevance in affective scenes is largely stable across perceivers.

Our findings extend prior work by directly linking social features in visual scenes to affective valence and arousal, demonstrating that social and emotional appraisals are not independent but co-organized into structured, interpretable dimensions. Emotion sharing, action coordination, and perceived equality consistently covaried with pleasantness ratings and were robust across observers, reinforcing the idea that social meaning constitutes a core, rather than peripheral, dimension of affective scene perception. This structured co-variation offers a more nuanced account of how individuals evaluate complex scenes—one that incorporates interpersonal context as an integral component of emotional appraisal. By providing a validated and openly accessible dataset annotated along multiple social-affective dimensions, our study also addresses a practical need for standardized tools that enable targeted stimulus selection in experimental research. Together, these contributions support both theoretical refinement and methodological precision in the study of social relevance.

These findings have direct implications for the design of future experiments in affective neuroscience, social cognition, and related fields. Rather than relying on coarse categorical contrasts—such as the mere presence or absence of human figures—researchers should consider selecting stimuli based on multidimensional social-affective properties. For example, studies on empathy, moral judgment, or interpersonal decision-making may benefit from using stimuli that systematically vary in engagement-related aspects, such as action sharing or perceived equality. Additionally, incorporating objective image features – such as number of individuals depicted, face visibility, or gaze direction – can facilitate hybrid models that integrate low-level perceptual input with high-level social meaning. Our dataset provides a structured foundation for such approaches, supporting more ecologically valid and theoretically precise investigations of socio-affective information processing.

Several limitations should be acknowledged. First, although our stimuli spanned a range of emotional and social content, all ratings were based on static images presented without contextual or temporal information. While this format is widely used in affective neuroscience, it may limit ecological validity and underrepresent the dynamic nature of social interactions. Nonetheless, by working within the constraints of a standardized and widely adopted image set, we provide an empirically grounded resource for improving stimulus selection. Second, the final stimulus set was not fully balanced across all dimensions of interest, such as interaction type, valence, and arousal. As a result, some combinations of features, e.g., negative group scenes with high equality, were underrepresented due to natural constraints in the IAPS database, potentially limiting the generalizability of fine-grained comparisons. Finally, while our rating dimensions were both theory-driven and empirically supported, they do not exhaust the full range of social meaning. Future extensions might incorporate additional constructs such as perceived affiliation, threat, or intentionality to further refine the structure of social-affective appraisal.

Taken together, our study provides a theoretically informed, methodologically rigorous framework for quantifying social relevance in visual scene perception. By bridging the gap between standardized affective image sets and the complex interpersonal appraisals that guide human social behavior, we offer a novel resource for investigating how emotional and social meanings co-construct perceptual experience. As research increasingly emphasizes the relational structure of social cognition—including spatial, hierarchical, and affective dimensions—our work contributes an accessible and empirically validated toolset for operationalizing these features in experimental designs. Future research may use our dataset to test mechanistic models of social relevance computation, explore developmental or cultural differences in social appraisal, or integrate behavioral and neural measures to advance comprehensive accounts of how humans perceive, interpret, and respond to social meaning in complex environments.

## Acknowledgements

This study was funded by the Deutsche Forschungsgemeinschaft (DFG, Project-ID 454648639 - SFB 1528). The code of this study is available upon request from the corresponding author AS. Data and analyses code are available on the Open Science Framework (https://osf.io/9m3ef/?view_only=082eceb798aa4cf88792657ca1f03cb1). We thank Roger Mundry for making functions for the analyses available and Weicheng Lian for assistance in data collection.

## CRediT Authorship Contribution Statement

**Vanessa Mitschke:** Conceptualization, Methodology, Project administration, Writing – original draft, Writing – review & editing. **Francesco Grassi:** Data curation, Formal analysis, Visualization, Writing – original draft, Writing – review & editing. **Alperen Doganer:** Writing – review & editing. **Anne Schacht:** Conceptualization, Funding acquisition, Methodology, Project administration, Resources, Supervision, Validation, Writing – original draft, Writing – review & editing.

## Supplementary Material

### S1. Results of the Exploratory Correlation Analysis

**Table S1.** Semimatrix of d-correlations between ratings and objective features for single-person images.

|  | Pleasantness | Participation | Face<br>Visibility | Gaze Direction<br>Viewer | N Faces |
| --- | --- | --- | --- | --- | --- |
| Social Relevance | 0.95 | 0.29 | 0.18 | -0.01 | 0.36 |
| Pleasantness |  | 0.31 | 0.14 | 0.00 | 0.29 |
| Participation |  |  | 0.06 | 0.01 | 0.01 |
| Face Visibility |  |  |  | -0.02 | 0.77 |
| Gaze Direction<br>Viewer |  |  |  |  | -0.01 |

**Table S2.** Semimatrix of d-correlations between ratings and objective features for multiple-person images.

|  | Pleasantness | Participation | Emotion<br>Sharing | Action<br>Sharing | Equality | Face<br>Visibility | Gaze Direction<br>Viewer | Gaze Direction<br>Within | N<br>Faces | N<br>People | People<br>Group |
| --- | --- | --- | --- | --- | --- | --- | --- | --- | --- | --- | --- |
| Social Relevance | 0.92 | -0.09 | 0.78 | 0.76 | 0.70 | 0.04 | -0.02 | 0.04 | 0.17 | 0.13 | 0.00 |
| Pleasantness |  | -0.04 | 0.76 | 0.70 | 0.70 | 0.04 | 0.00 | 0.02 | 0.11 | 0.07 | 0.00 |
| Participation |  |  | -0.06 | -0.02 | -0.11 | 0.02 | -0.02 | -0.03 | 0.09 | 0.30 | 0.06 |
| Emotion<br>Sharing |  |  |  | 0.95 | 0.90 | 0.07 | -0.05 | 0.05 | 0.2 | 0.18 | 0.01 |
| Action Sharing |  |  |  |  | 0.81 | 0.08 | -0.04 | 0.03 | 0.21 | 0.21 | 0.01 |
| Equality |  |  |  |  |  | 0.05 | -0.04 | 0.07 | 0.21 | 0.17 | 0.00 |
| Face Visibility |  |  |  |  |  |  | 0.05 | -0.03 | 0.38 | -0.02 | 0.10 |
| Gaze Direction<br>Viewer |  |  |  |  |  |  |  | -0.10 | 0.17 | 0.05 | 0.01 |
| Gaze Direction<br>Within |  |  |  |  |  |  |  |  | -0.05 | 0.02 | 0.00 |
| N Face |  |  |  |  |  |  |  |  |  | 0.55 | 0.13 |
| N People |  |  |  |  |  |  |  |  |  |  | 0.41 |

### S2. Factor Analysis on Ratings of Engagement

Given the very strong correlation between emotion sharing, action sharing and equality, we conducted an Exploratory Factor Analysis (EFA) to investigate how well these three variables load onto a potential underlying latent factor. The analysis was conducted in R using function *fa* from package *psych* (version 2.5.3, Revelle, 2007).

Since we conducted EFA on three variables, we could only extract one latent factor. In fact, degrees of freedom in an EFA are calculated as:

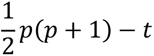

Where 𝑝 is the number of observed variables and 𝑡 the number of parameters being estimated (including both factor loadings and the unique variance). Calculating one factor from three observed variables means estimating three factor loadings and three respective unique variances (𝑡 = 6). Therefore, we end up with **zero degrees of freedom.** Trying to calculate even more factors would end up estimating more parameters than the data can support.

Please note that a consequence of having zero degrees of freedom, is that the model estimated here is **saturated**. The estimated factor will explain the shared variance among variables, but **there is no way to statistically test** if this factor solution is a good fit or not. Therefore, here we won’t report measures of model fit.

The results of the EFA are showed in (**Table S3**).

**Table S3.** Results of the EFA on the ratings of emotion sharing, action sharing and equality.

| Rating Variable | Factor 1 | Communality | Uniqueness | Complexity |
| --- | --- | --- | --- | --- |
| Emotion Sharing | 1.00 | 1.00 | 0.00 | 1 |
| Action Sharing | 0.95 | 0.91 | 0.09 | 1 |
| Equality | 0.91 | 0.82 | 0.18 | 1 |

The EFA revealed that emotion sharing, action sharing and equality load quite clearly on a single factor explains a **very high portion of total variance** (91%). **A**ll three variables show **very large positive loadings** on the factor (> 0.90). **Compared to the other two variables, equality** contributes a slightly **less** to the **communal variance** (h^2^ = 0.82) and has a slightly **larger uniqueness** (u^2^ = 0.18).

### S3. Interaction Effect of Scene Participation and Person Number of Social Relevance Rating

The LMM used to test the effect of scene pleasantness on social relevance, while controlling for scene participation, person number and gaze direction also showed a significant interaction effect of participation and person number (*β* = -0.16, *SE* = 0.03, *p* < .001, 95% CI [-0.22, -0.11], see **Table 5** in the main text). For single-person images, increased ratings of scene participation resulted in increased social relevance, while this effect was drastically reduced for multiple-person images (**Figure S1**).

**Figure S1.**
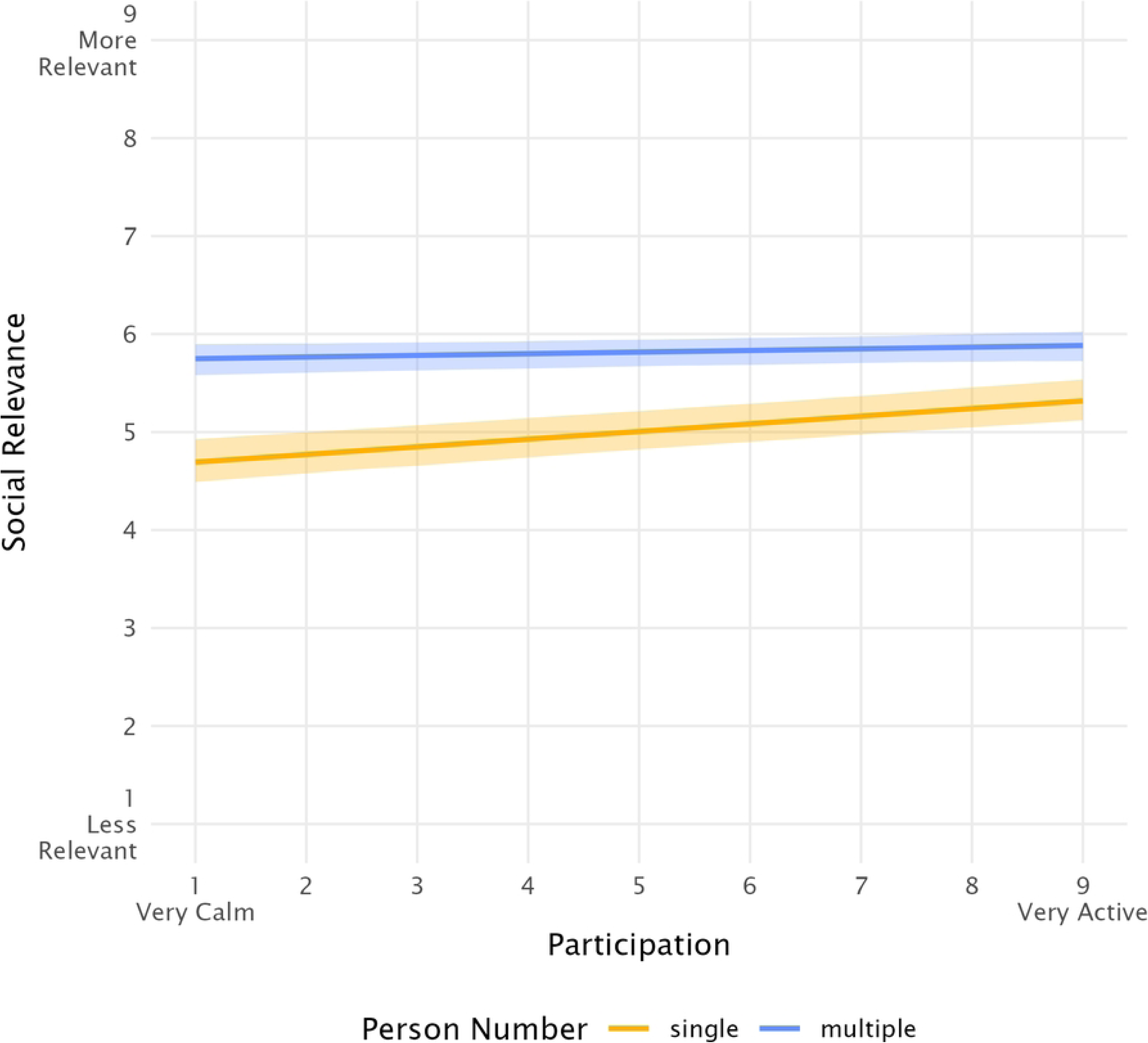
Social relevance as a function of scene participation, for single- and multiple-person images. Shaded areas represent 95% confidence intervals.

### S4. Main Effects of Scene Participation and Gaze direction of Social Relevance Rating

The LMM used to test the effect of engagement on social relevance, while controlling for scene participation and gaze direction (**Table 6** in the main text) also showed a significant main effect of scene participation (*β* = 0.10, *SE* = 0.03, *p* < .001, 95% CI [0.05, 0.15]), with increased participation resulting in increased social relevance, and a main effect of gaze direction (*β* = -0.84, *SE* = 0.35, *p* = .017, 95% CI [-1.54, -0.11]), with lower ratings of social relevance for images with averted compared to direct gaze (**Figure S2**).

**Figure S2.**
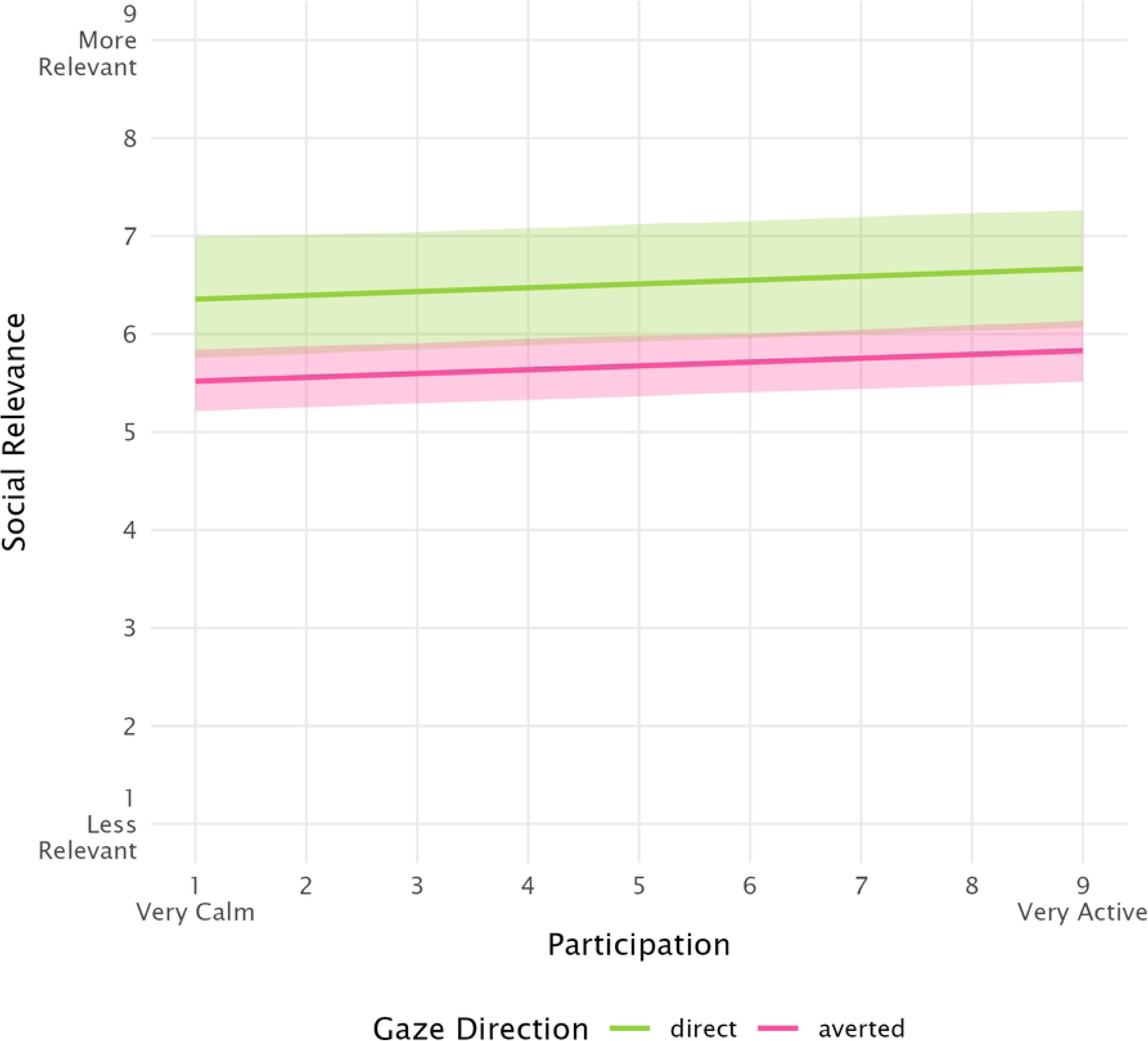
Social relevance as a function of scene participation, for direct and averted gaze direction. Shaded areas represent 95% confidence intervals.

## Notes

### Competing Interest Statement

The authors have declared no competing interest.

